# Global expression profile of Enterohemorrhagic *Escherichia coli* O157:H7 in phagosome of murine macrophages

**DOI:** 10.1101/2024.12.18.629128

**Authors:** Nahuel Agustín Riviere, Lin Jiang, Xiuju Wu, María Carolina Casabonne, Lu Guo, Libia Yael Smith, Wanderson Marques Da Silva, Ángel Adrián Cataldi, Ke Xing, Jinlong Bei, Mariano Larzábal

## Abstract

EHEC O157:H7 is responsible for Hemolytic Uremic Syndrome (HUS) outbreaks in humans and its virulence is associated with Shiga toxin (Stx). During the pathogenesis of HUS, EHEC O157:H7 is phagocytized by the intestinal host macrophage into the phagosome. According to our study, a low percentage of phagocytized bacteria survived in the murine macrophage phagosome. To understand the initials mechanisms involved in bacterial persistence, we performed a global profile of bacterial RNAseq analysis in the early phagosome of murine macrophage and *in vitro* assays with more extreme stress conditions. The defense strategy of an early murine phagosome consisted on producing DNA damage, membrane damage, acid pH and nitric oxide (NO) agents. These hostile conditions triggering the bacterial SOS response, lipid biosynthesis and NO detoxification enzymes in response to DNA damage, membrane damage and nitric oxide agents, respectively. In turn, to survive, the bacterium conserves energy by downregulating flagellar biosynthesis, T3SS and T6SS virulence mechanisms. In contrast, *stx2a* expression is upregulated. At the same time, it increases the ribosomal levels and amino acid synthesis to react more effectively against adverse conditions. Under more extreme stress conditions, EHEC O157:H7 expressed genes related to an acidic environment (pH 4.5) upregulating the acid stress response pathways AR2-5, whereas, upon high concentrations of hydrogen peroxide, it transcribed higher levels of genes of the OxyR pathway and subsequently hydrogen peroxide oxidative stress-related genes.

## Introduction

Enterohemorrhagic *Escherichia coli* (EHEC) is a foodborne pathogen of worldwide importance associated with bloody diarrhea and Hemolytic Uremic Syndrome (HUS) outbreaks in humans [1], whose reservoirs are ruminants, mainly bovines [2]. Consumption of contaminated food or water is the primary source of human infection [3]. EHEC O157:H7 is a serotype responsible for the majority of the most severe cases of diarrheal diseases [4]. The main virulence factor of EHEC O157:H7 is the Shiga toxin (*stx*) encoded by lysogenic lambdoid bacteriophage [5]. EHEC O157:H7 EDL933 strain encodes Stx_1_ and Stx2a. While Stx_1_ has been connected to human illnesses, Stx2a subtypes have shown a stronger correlation with the onset of HUS [6]. EHEC O157:H7 also has a Type 3 Secretion System (T3SS), a virulence mechanism that is encoded in the pathogenicity island named Locus of Enterocyte Effacement (LEE) [7]. During the infection process, EHEC moves through the gastrointestinal tract, where the bacteria survive the acidic condition of the stomach. T3SS expression is essential for intestinal colonization and is responsible for the characteristic “attaching and effacing” lesion [8]. The intestinal adhesion occurs exclusively in regions with overlying follicle-associated epithelium of Peyer’s patches [9]. These follicles contain M cells responsible for translocating pathogens from the intestinal lumen to the basolateral side of the epithelium, where the bacteria are then phagocytized by macrophages [9]. In the case of EHEC O157:H7, the macrophage internalizes this bacterium into the early phagosome, which then undergoes maturation [10]. Phagosome acidification begins after phagocytosis, reaching pH values close to 4.5 [11].

During phagocytosis of microorganisms, oxygen is reduced to O_2_^-^ via NADPH oxidase, which increases reactive oxygen species (ROS) levels, an essential antimicrobial mechanism of phagocytes that restricts oxygen in solution, thus generating a hypoxic and oxidative environment [12]. Another defense strategy by the macrophage is limiting access to essential nutrients, such as carbon sources and metal ions like iron, manganese, magnesium, copper, and zinc [13]. EHEC O157:H7 phagocytosis subsequently triggers a bacterial SOS response because of the DNA damage caused by early phagosome stress conditions [14]. This response induces the lytic state of lambda bacteriophage and the subsequent Stx expression, which provokes cytotoxicity in the macrophage [15]. However, limited information is available regarding the interactions of the EHEC O157:H7 strain with host immune cells.

Bacterial transcriptome expression analysis (RNAseq) has become a fundamental tool for understanding molecular changes during pathogenesis. In this work, we decided to deepen the knowledge of the mechanisms involved in the response of EHEC O157:H7 strain inside the early phagosome. Moreover, we decided to simulate some of the biochemical properties of a late macrophage phagosome *in vitro* to study the bacterial transcriptome under these conditions.

## Material and methods

### Bacterial strain, macrophages cells and growth conditions

EHEC O157:H7 strain EDL933 was isolated from ground beef in 1982, during an EHEC outbreak in the United States [16]. The EHEC O157:H7 EDL933 strain was sequenced and deposited at NCBI (CP008957.1). EHEC O157:H7 EDL933 strain encodes 5920 genes in a 5.5 Mb genome 50,5 % GC and contains a megaplasmid (pO157) of 94.7 kb with approximately 136 genes [17]. The functional studies were performed by growing the bacteria strain in Luria-Bertani (LB) medium overnight at 37°C and 200 g; then, the bacteria were diluted 1/50 in LB fresh and grown to exponential phase (optical density (OD) at 600 nm of 0.6), at 37°C and 200 g.

The murine macrophage cell line used for the EHEC O157:H7 EDL933 virulence assays was RAW 264.7. This cell line established from a tumor in a male mouse induced by the Abelson murine leukemia virus. RAW 264.7 was cultured in DMEM (Gibco, Thermo Fisher, USA) supplemented with 10% fetal bovine serum (FBS) at 37°C under a 5% CO_2_ atmosphere.

### Fluorescence Confocal Microscopy of phagocytized EHEC O157:H7

The intracellular survival of EHEC O157:H7 in RAW 264.7 murine macrophages after 24 h of incubation was assessed by confocal microscopy. For this purpose, the RAW 264.7 cells were grown on cover slips in a 12-well plate until 90% confluency (1.10^6^ cells per well). In parallel, a grown culture of EHEC O157:H7 was incubated with Carboxyfluorescein succinimidyl ester (CFSE) for 30 min. Subsequently, the culture was incubated with the macrophage monolayer at an MOI of 100. After 24 h of incubation at 37°C and 5% CO_2_, the monolayer was washed three times with PBS before adding 200 µl/well of Lysotracker red (Termofisher, USA; diluted 1:1000 in PBS) to perform the labelling with an incubation for 1 h at 37°C in the dark.

After the incubation, the EHEC-incubated macrophage monolayer was fixed with 4% paraformaldehyde (PFA) in PBS for 20 min. The monolayer was washed three times with PBS and the cells were permeabilized by adding first 200 µl/well of 0.15% Triton (incubation for 15 min) and subsequently 200 µl/well of 1% BSA to block and prevent non-specific labelling (incubation for 15 min). The labelling was carried out with the primary antibody LAMP3 (Lysosomal Associated Membrane Protein 3, Termofisher, USA) diluted in PBS 1X (1:50), with an incubation for 1.5 h. After three washes with PBS, a second incubation with the secondary antibody Cy5 (Termofisher, USA) diluted in PBS (1:600) was performed for 1.5 h. Finally, the coverslips were washed three times with PBS mounted on slides and sealed with VectaShield. The samples were analyzed by confocal microscope-Leica TCS SP5-CICVyA-INTA.

### EHEC survival assay in RAW 264.7 murine macrophages

RAW 264.7 cells were seeded at 1.10^6^ per well in a 12-well plate and co-incubated with EHEC O157:H7 bacterial culture at an OD_600_ of 0.6 and at an MOI of 100 (1.10^8^ CFU). The infections were synchronized by incubating the plates for 30 min (time 0h) at 37°C and 5% CO_2_ after a centrifugation at 400 g for 5 min. Subsequently, the infected cells were washed three times with PBS. Then, DMEM medium supplemented with 10% SFB and 100 µg/ml gentamicin was added to the monolayer to eliminate non-phagocytized extracellular bacteria. After 2 h of incubation at 37°C and 5% CO_2_, the infected cells were washed three times with PBS and lysed with 0.025% SDS solution. Serial dilutions were then performed in PBS and plated on LB agar to quantify the phagocytized bacteria (CFU/ml).

The assessment of the intracellular survival of EHEC O157:H7 in macrophages was performed by replacing the DMEM medium supplemented with 10% SFB and 100 µg/ml gentamicin with DMEM 10% SFB containing 15 µg/ml gentamicin. In parallel, cell cultures were analyzed for viable bacteria after 24 h incubation at 37°C under 5% CO_2_. Subsequently, the infected cells were washed three times with PBS before performing lysis with 0.025% SDS solution. Serial dilutions of the lysates in PBS were seeded on LB agar to quantify the surviving bacteria (CFU/ml). The percentage of surviving bacteria was relativized to the number of bacteria phagocytized after 2 h.

### EHEC under *in vitro* stress conditions

EHEC O157:H7 was exposed to *in vitro* stress conditions to assess RNA transcription levels. The EHEC O157:H7 culture was grown on LB medium to OD_600_ 0.6 and centrifuged at 4000 g for 5 min. Then, the bacteria were washed three times with PBS before adding M9 minimal medium, a medium with no added carbon sources, to recreate a nutrient-free environment [18,19]. The medium was supplemented with 800 mM hydrogen peroxide, to recreate oxidative stress, and 400 µM bipyridil, as a divalent cation chelator. The growth condition was at pH 4.5 to generate acid stress and low oxygen tension. This condition was achieved by covering the tube with the culture minimal medium and without shaking to avoid aeration. Total RNA was extracted from cultures incubated for 2 h under the described conditions.

### RNA-Seq expression profiling: RNA isolation

A transcriptomic assay was performed using the RNA-seq methodology, which allowed obtaining a gene expression profile of EHEC O157:H7 EDL933 strain phagocytized by RAW 264.7 murine macrophages or under stress conditions at 2 h post-incubation.

For the phagocytized EHEC O157:H7 assays, three biological replicates were made; each replicate consisted of six 12-well plates. Each well had 1.10^6^ RAW 264.7 cells at close to 90% confluence. The RAW 264.7 cells murine macrophages were infected with a bacterial culture of EHEC with 1.10^8^ CFU to obtain an MOI of 100. At 2 h post infection, the infected macrophages were lysed in ice-cold ‘RNA stabilization solution’ (0.025% SDS, 19% ethanol, 1% acidic phenol in water) and incubated on ice for 30 min. The lysate containing EHEC O157:H7 was collected by centrifugation at 4000 g 4°C for 10 min. The supernatant was discarded, and the pellet was washed three times with a phenol-alcohol solution (1%:19%, respectively). Then, the cell pellet was dissolved in 1 ml of TransZol (Transgen Biotech, China) on ice and 200 µl of chloroform. The samples were centrifuged at 12000 g, 4°C for 15 min.

Then, the aqueous phase was transferred and total RNA purification was carried out following the indications of the Monarch Total RNA Miniprep kit (New England Biolabs, USA).

The sample was enriched with bacterial RNA by performing a second RNA purification step with the MICROBEnrich kit (Ambion, USA) to eliminate the eukaryotic RNA. The kit employs magnetic beads derived with an oligonucleotide that hybridizes to capture polyadenilated 3‘ ends of eukaryotic mRNAs.

In the case of the assays of EHEC under stress conditions, the culture was centrifuged and the pellet was dissolved with 1 ml of TransZol (Transgen Biotech, China). Then, 200 µl of chloroform was added and the samples were centrifuged at 10000 g, 4°C for 15 min. Then, the aqueous phase was transferred to a new Eppendorf tube to perform total RNA purification according to the indications of the Monarch Total RNA Miniprep kit (New England Biolabs, USA). In this case, the bacterial RNA enrichment step with the microbenrich kit was dispensable, since no murine macrophage was used. Control RNA was isolated from bacterial cells grown in DMEM and M9 minimal medium.

The RNA integrity was verified in all samples using an Agilent Bioanalyzer 2100 and the concentration was measured using the Qubit fluorometer. See Supplementary Figure 1 for quality control and S1 Table for RNA concentrations.

### Library preparation and deep sequencing

Total RNA obtained from biological replicates were initially treated with Ribozero (Illumina inc., USA) to remove ribosomal RNA (rRNA). The removal of rRNA in the EHEC O157:H7 RNA samples obtained from the RAW 264.7 infection was improved by performing the RiboZero Plus protocol. The RNA samples of the controls and of EHEC exposed to stress *in vitro* were processed with the RiboZero Bacterial protocol (Illumina inc, USA.). Subsequently, the samples were quantified with Qubit (Termofisher, USA), and analyzed with Fragment Bioanalyzer (Agilent, USA).

The construction of the cDNA libraries with single-ended ends for 300 bp fragments was performed following the Illumina Truseq protocol (Illumina inc. USA). The cDNAs were purified by beads and analyzed by capillary electrophoresis in Agilent Bioanalyzer 2100 (Agilent Technologies, USA). The sequencing was performed on the Illumina MiSeq platform (Illumina inc. USA). To achieve a higher sequencing depth, we generated about 70 million reads for phagocytized EHEC and 10 million reads for EHEC subjected to stress conditions.

### Bioinformatics analyses

The raw data obtained from RNA sequencing was subjected to quality examination by using FastQC v0.11.9 Babraham Bioinformatics - FastQC A Quality Control tool for High Throughput Sequence Data, n.d.). Trim Galore v.0.6.10 (https://github.com/FelixKrueger/TrimGalore) and Cutadapt v.3.4 [21] were subsequently used to generate high quality clean reads by removing sequences containing more than 5% of N base, adapter sequences, and reads with more than 10% *Q* < 20 bases. After preprocessing, 10,593,521∼ 76,682,018 clean reads were finally obtained for each sample.

Gene expression was quantified by mapping the clean reads to the reference genome (CP008957.1) by using HISAT2 v2.2.1 [22]. Stringtie v2.2.0 [23] was employed for quantitative analysis. The analysis of expression differences was then performed in R package DEseq2 [24], with a model based on the negative binomial distribution. Differentially Expressed Genes (DEGs) was defined according to the criterion: log_2_ fold change >1 or log_2_ fold change <-1 & *fdr* <0.05. Enrichment analyses of DEGs against Gene Ontology and KEGG database were performed with R package clusterProfiler v4.0 [25].

## Results and discussion

### Phagocytosis levels and survival of EHEC O157:H7 strain in murine macrophages

Using fluorescence confocal microscopy, EHEC O157:H7 was visualized within macrophages, indicating the pathogen’s ability to persist in this cellular environment 24 h post-incubation (Fig 1A). Colocalization between acidic compartments (blue dots) and EHEC O157:H7 (green) was observed (Fig 1A).

**Figure 1AB.**
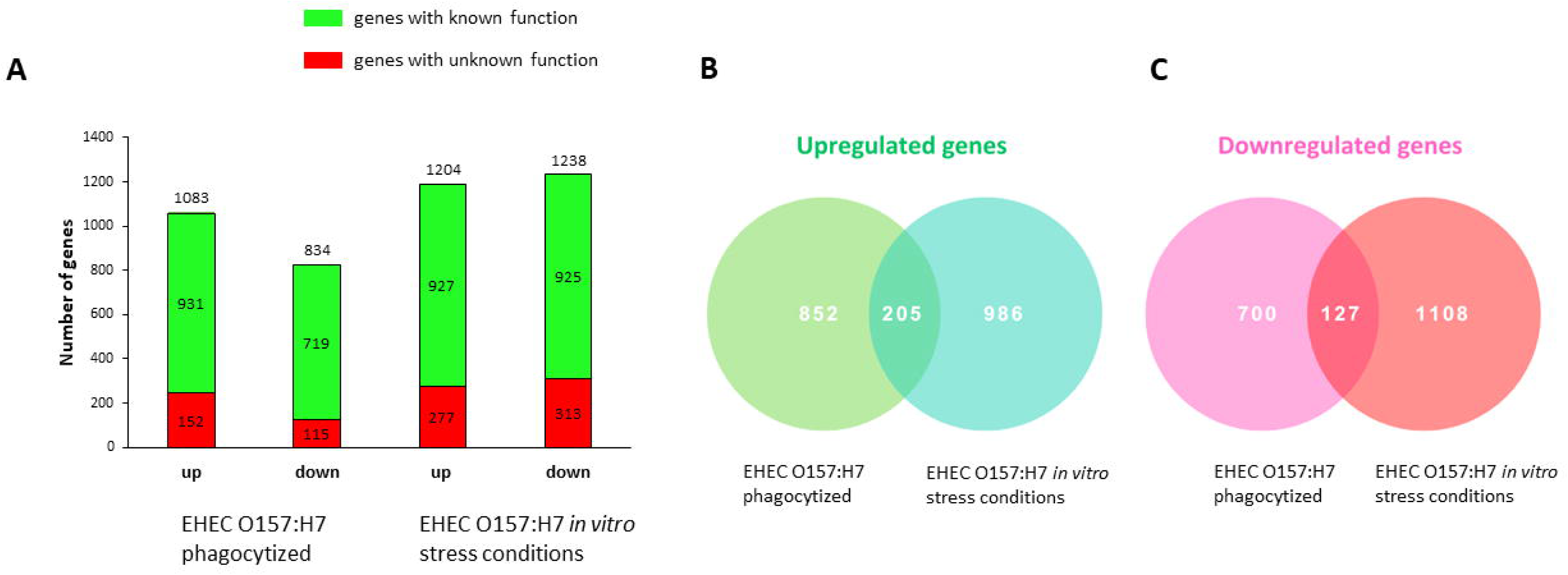
Infection of murine macrophages with EHEC O157:H7. A) Images obtained by confocal microscopy of macrophages incubated with EHEC O157:H7 for 24 h. The macrophage monolayer was observed by differential interference contrast (DIC). Macrophages were labeled with LAMP3 (Lysosome-associated membrane glycoprotein 3) (blue); Prior to infection, the bacteria were labeled with CFSE (carboxyfluorescein succinimidyl ester, Termofisher, USA) (green), a molecule that emits fluorescence when intracellular esterases in living cells cleave the acetate group, generating a green fluorescent carboxyfluorescein molecule, which becomes impermeable. The layers were superimposed (merge) and EHEC O157:H7 could be observed inside the macrophage. B) Intracellular survival of EHEC O157:H7 in RAW 264.7 macrophages. RAW 264.7 cells were incubated with EHEC O157:H7 at an MOI of 100 for 30 min. Then, the medium was replaced with DMEM containing 10% SFB and 100 µg/µl gentamicin for 2 h to kill extracellular bacteria. The cells were then incubated for 24 h with DMEM containing 15 µg/µl gentamicin at 37°C under 5% CO_2_. Lysates were then plated to count viable intracellular bacteria. The percent bacterial survival was calculated based on viable counts (CFU/ml) relative to 0 h. The errors bars represent SD from at least three independent experiments. *, P< 0,001, Unpaired t-test analysis.

Murine macrophage RAW 264.7 monolayers infected with EHEC O157:H7 strain phagocytized 0.1% (1.14×10^5^ CFU/ml) of the initial MOI at 2 h post infection (p.i.) (Fig 1B). Only 0.01% (2.3×10^3^ CFU/ml) of the initial infection bacteria persisted in the macrophage 24 h p.i. (Fig 1A and 1B). The obtained percent persistence indicated that the bacteria did not replicate within the phagosome. These results imply that, despite the low rate of phagocytosis and persistence, several elements expressed by EHEC O157:H7 strains may have the ability to influence survival during the process of phagocytosis.

### Transcriptomic profile of EHEC O157:H7 strain phagocytized by murine macrophages

We implemented a massive sequence by RNA-seq methodology to determine the transcriptomic profile of EHEC O157:H7 strain in response to early macrophage phagocytosis and in extreme stress conditions (Fig 2AB). Obtaining high-quality RNA that represented bacterial gene expression in interaction with the host was challenging. Some of the factors affecting the procedure included the low intake of phagocytic bacteria, the stability of the pathogen against phagosomés hostile environment, overexpression of host messenger RNA (less than 1% of bacterial RNA in sample), short half-life of RNA, contaminated sample with ribosomal RNA and small RNA. In parallel, we simulated some of the biochemical properties of a late macrophage phagosome *in vitro* to study the bacterial transcriptome under more extreme conditions.

**Figure 2AB.**
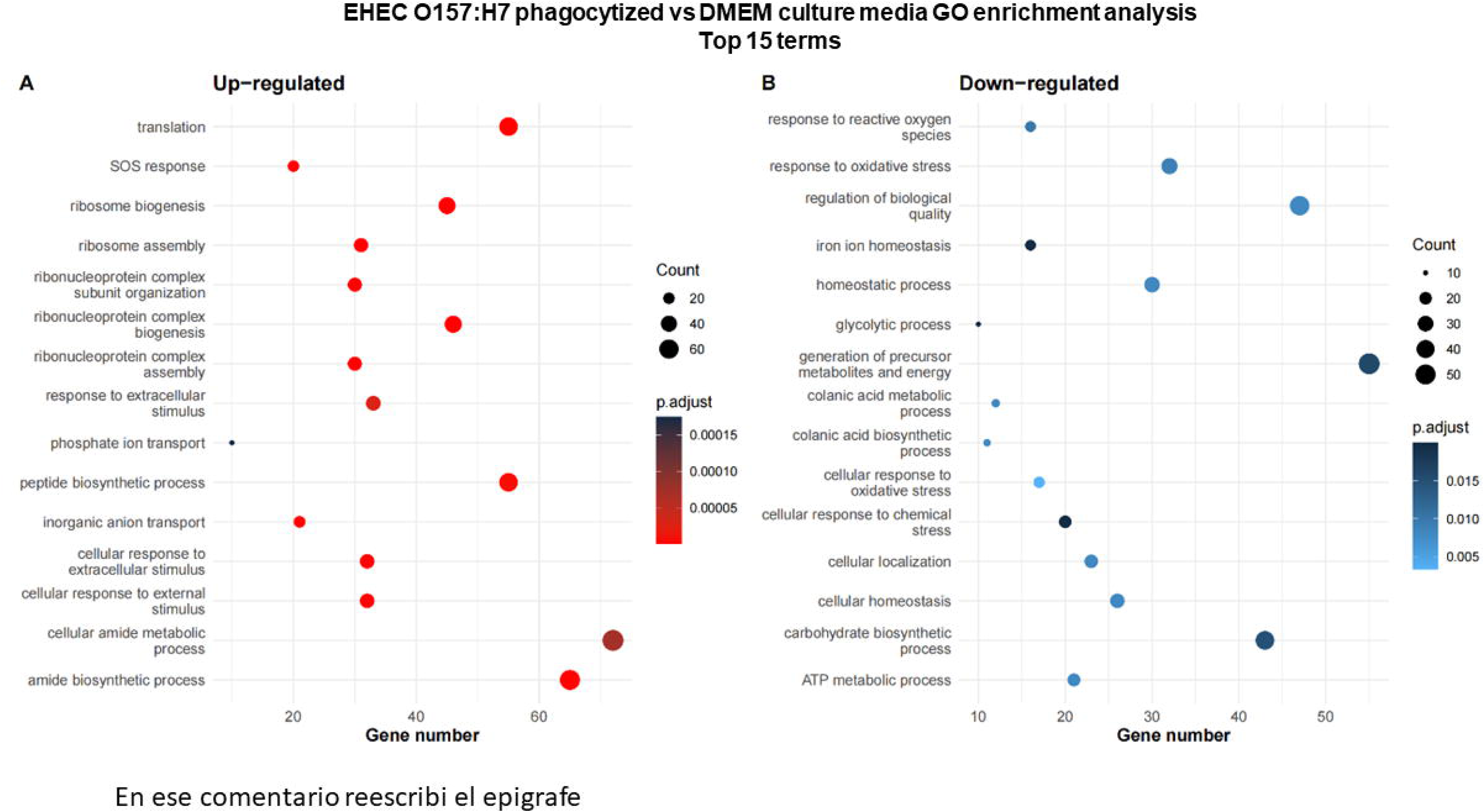
RNA-seq-based strategy. A) Phagocytized EHEC O157:H7. A fresh culture of EHEC O157:H7 was incubated with macrophages at an MOI of 100 for 2 h with gentamicin (Gentamicin Protection Assay). The phagocytized bacteria were recovered by differential lysis. Then, the RNA was extracted with TransZol (Materials and methods). The generated cDNA was sequenced for differential expression analysis and the data were compared against EHEC grown in DMEM medium (control). B) EHEC O157:H7 in stress conditions. In stress conditions, EHEC O157:H7 was subjected to a 2 h incubation period in M9 minimal medium, designed to simulate a late phagosome environment. Subsequently, RNA extraction was performed using the TransZol method (as detailed in the Materials and Methods section). The resulting cDNA was subjected to sequencing for differential expression analysis, and the obtained data were then compared to the gene expression profiles of EHEC grown under control conditions in M9 minimal medium, without any stressors.

The transcriptomic analysis of phagocytized EHEC O157:H7 strain by murine macrophage for 2 h allowed the detection of the expression of 5897 genes with a coverage of 99% according to GenBank annotation (CP008957.1) [16]. The comparison with the RNA obtained from bacteria grown in DMEM media (control) revealed that 1917 of all genes (32,65%) were differentially expressed: 1083 genes (18%) upregulated and 834 genes (14,65%) downregulated (S2 table). In addition, 152 (2,5%) of the upregulated genes and 115 (1,94 %) of the downregulated genes remained as genes of unknown function (Fig 3A).

**Figure 3ABC.**
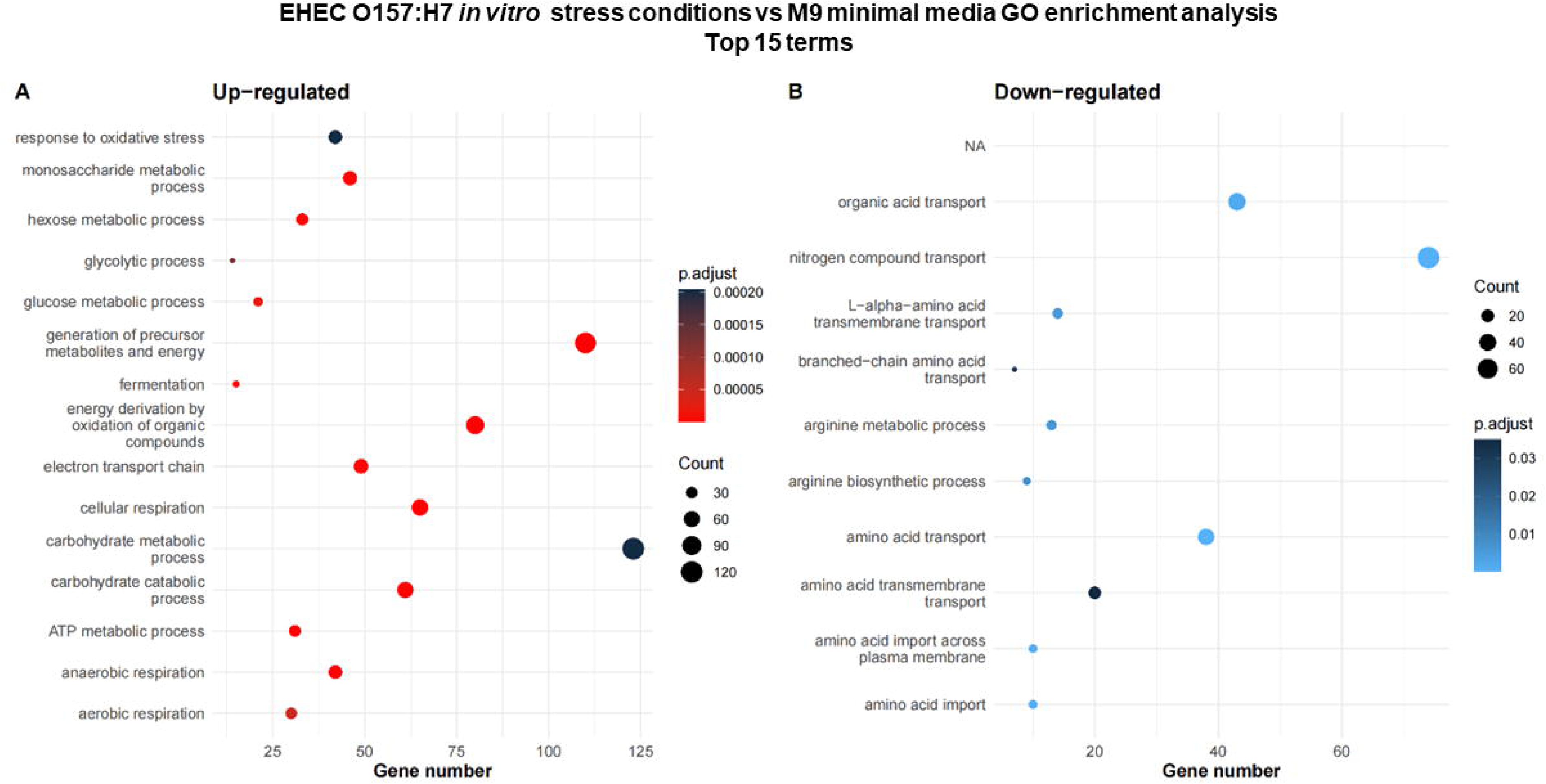
Global analysis of differentially expressed genes. A) Number of upregulated or downregulated genes for phagocytized and *in vitro* stress conditions EHEC O157:H7 at 2 h of incubation. Venn diagrams showing the number of genes upregulated B) or downregulated C) genes for phagocytized and *in vitro* stress conditions EHEC O157:H7 at 2 hs of incubation.

### Transcriptomic profile of EHEC O157:H7 strain exposed to *in vitro* stress conditions

The low rates of persisting bacteria in late stages of phagocytosis did not provide high-quality RNA for transcriptomics assays. Therefore, we simulated some of the biochemical properties of a late macrophage phagosome *in vitro* to study the bacterial transcriptome according to more extreme conditions (Fig 2B). Some of these biochemical conditions were consisted of an acid and oxidative environment with limited nutrients and divalent cations at 37°C.

Regarding the EHEC O157:H7 exposed to stress, were detected 5897 genes approximately a 99% of the genes present in the strain according to GenBank annotation (CP008957.1). Of these genes, 2442 (37 %) were differentially expressed: 1204 genes (20 %) upregulated and 1238 genes (20,9%) downregulated) in relation to the control condition (S2 Table). Around 277 (4,6%) of the upregulated genes and 313 (5,2%) of the downregulated genes had unknown function (Fig 3A).

The principal component analysis (PCA) revealed that samples of phagocytized EHEC O157:H7 and exposed to *in vitro* stress conditions clustered distinctly from the control samples along principal component 1 (PC1), indicating significant differences in their expression profiles. The clear separation along PC1 underscores the impact of the stress conditions on the bacterial response, suggesting that the experimental treatment induced a distinct and measurable shift in the bacterial gene expression. Additionally, the control samples exhibited some variability along principal component 2 (PC2), which could reflect inherent biological differences or experimental noise, but the overall clustering further supports the robustness of the observed effects (S2 Fig).

RNA-Seq is a highly robust and reliable technique for quantifying gene expression, offering greater sensitivity and accuracy compared to methods like microarrays. It sequences the entire transcriptome, providing reproducible, high-quality results that minimize the need for additional validation with qPCR. Advances in bioinformatics allow for stringent quality controls, ensuring the reliability of the data. In many cases, RNA-Seq data shows a strong correlation with qPCR, making additional validation redundant when the results have undergone necessary quality checks.

### Global expression analysis

According to a Venn diagram performed to determine common differential expression patterns, 852 upregulated genes were unique to the phagocytized bacteria, whereas 986 were exclusive of the *in vitro* stress condition (Fig 3B). In addition, 205 differentially expressed genes showed an upregulated profile in both cases. On the other hand, both conditions yielded quantities of unique downregulated genes (700 for the phagocytized bacteria; 1108 for the *in vitro* stress condition) and shared 127 downregulated genes (Fig 3C).

### Enrichment analysis of pathways and GO terms for DEGs in EHEC O157:H7 phagocytized or exposed to *in vitro* stress conditions

The analysis of DEGs of EHEC O157:H7 revealed that the top 15 upregulated biological enrichment profiles of phagocytized EHEC O157:H7 responded to proteins related to translation, SOS response, ribosome biogenesis, and peptide metabolism, among others, regarding gene ontology (GO) classification (Fig 4A). On the other hand, the downregulated genes included genes of the biological processes associated with generation of precursor metabolites and energy, carbohydrate biosynthetic process, regulation of biological quality and response to oxidative stress (Fig 4B).

**Figure 4AB.**
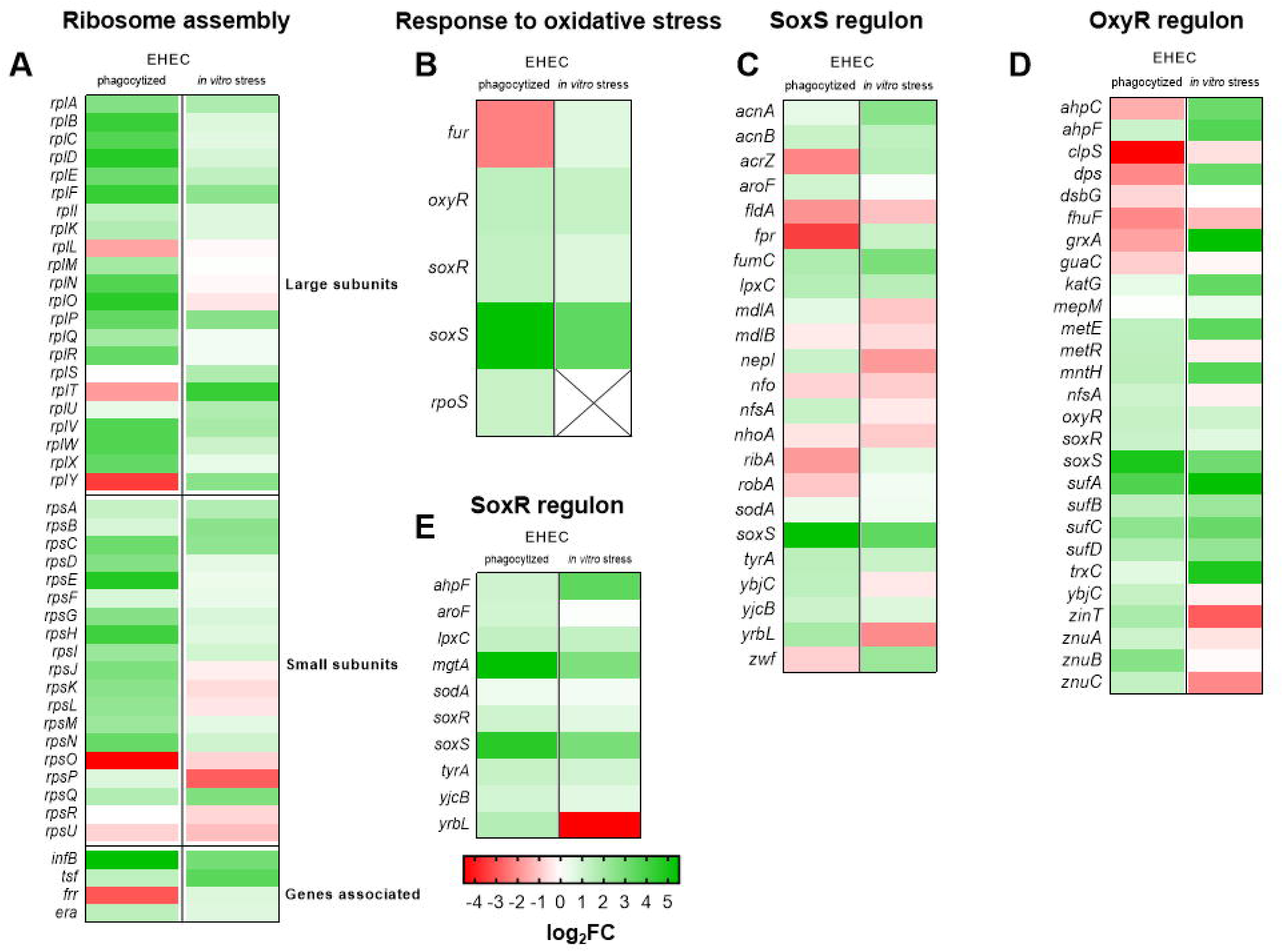
GO enrichment analysis of processes modulated in phagocytized EHEC O157:H7. The analysis was applied to identify GO terms from the domain “Biological processes” enriched for either upregulated (panel A; log_2_FC>1) or downregulated (panel B, log_2_FC<-1) differentially expressed genes at 2 h p.i. The size of the points represents the number of genes and the color refers to the statistical significance depending on the padj.

Regarding the genes expressed by EHEC O157:H7 grown under stress conditions, the top 15 upregulated biological enrichment profiles were associated with the generation of precursor metabolites and energy, carbohydrate metabolic processes and response to oxidative stress (Fig 5A). In contrast, the expression of genes associated with transport of organic acids and nitrogenous compounds showed downregulated levels (Fig 5B).

**Figure 5AB.**
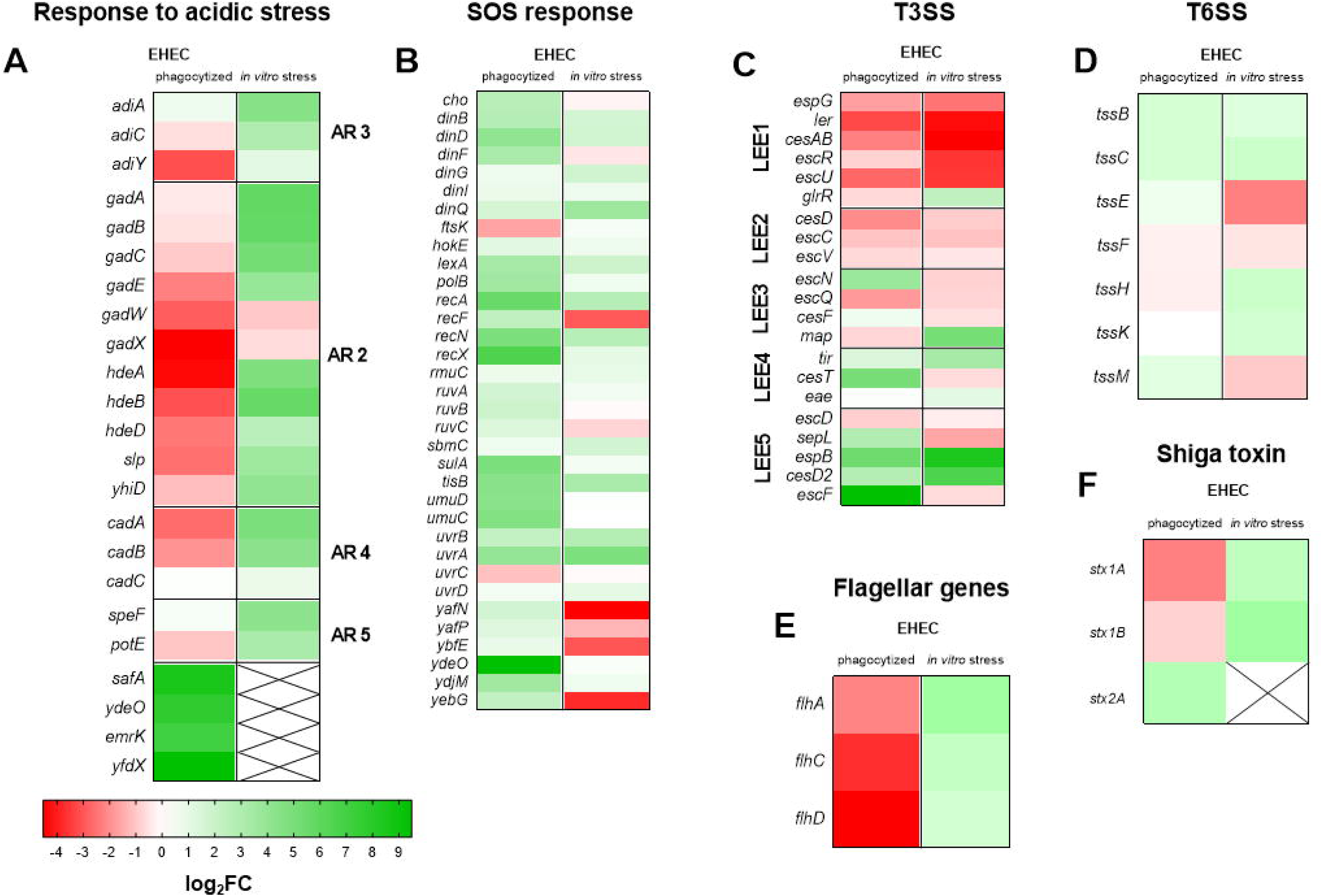
GO enrichment analysis of processes modulated in EHEC O157:H7 under stress conditions. Analysis was applied to identify GO terms from the domain “Biological processes” enriched for either upregulated (panel A; log_2_FC>1) or downregulated (panel B, log_2_FC<-1) differentially expressed genes at 2 h of incubation in minimal medium M9. The size of the points represents the number of genes and the color the statistical significance depending on the padj.

### Genes involved in ribosome assembly

Ribosome assembly is a tightly controlled process in which ribosomal proteins and subunits must interact and coordinate to promote the assembly of the mature particle [26]. Prokaryotes have 70S ribosomes, each consisting of a small (30S) and a large (50S) subunit. *E. coli*, for example, has a 16S RNA subunit (consisting of 1540 nucleotides) bound to 21 proteins. The large subunit consists of a 5S RNA subunit (120 nucleotides), a 23S RNA subunit (2900 nucleotides) and 31 proteins [27]. The analysis demonstrated that genes involved in ribosomal assembly displayed upregulated profiles in both stress conditions.

This group of genes are located in ribosomal operons. They include genes such as *rpla, rplv, rplf, rpma, rplp*, *rpmc* and *rpsc* (Fig 6A). Regarding the proteins encoded by these genes, all binds specifically to the 23S rRNA subunit, with an essential role during the early stages of 50S assembly [28], whereas the protein encoded by *rpsc* binds to the 30S subunit. Thus, the ribosomal biogenesis is associated with the need of a rapid response to the environmental changes occurring in the macrophage or during stress condition for bacteria to survive [29]. A severe reduction in translation could be detrimental during stress, precisely when bacteria need new protein synthesis to repair damage and adapt to the new environment.

**Figure 6ABCDE.**
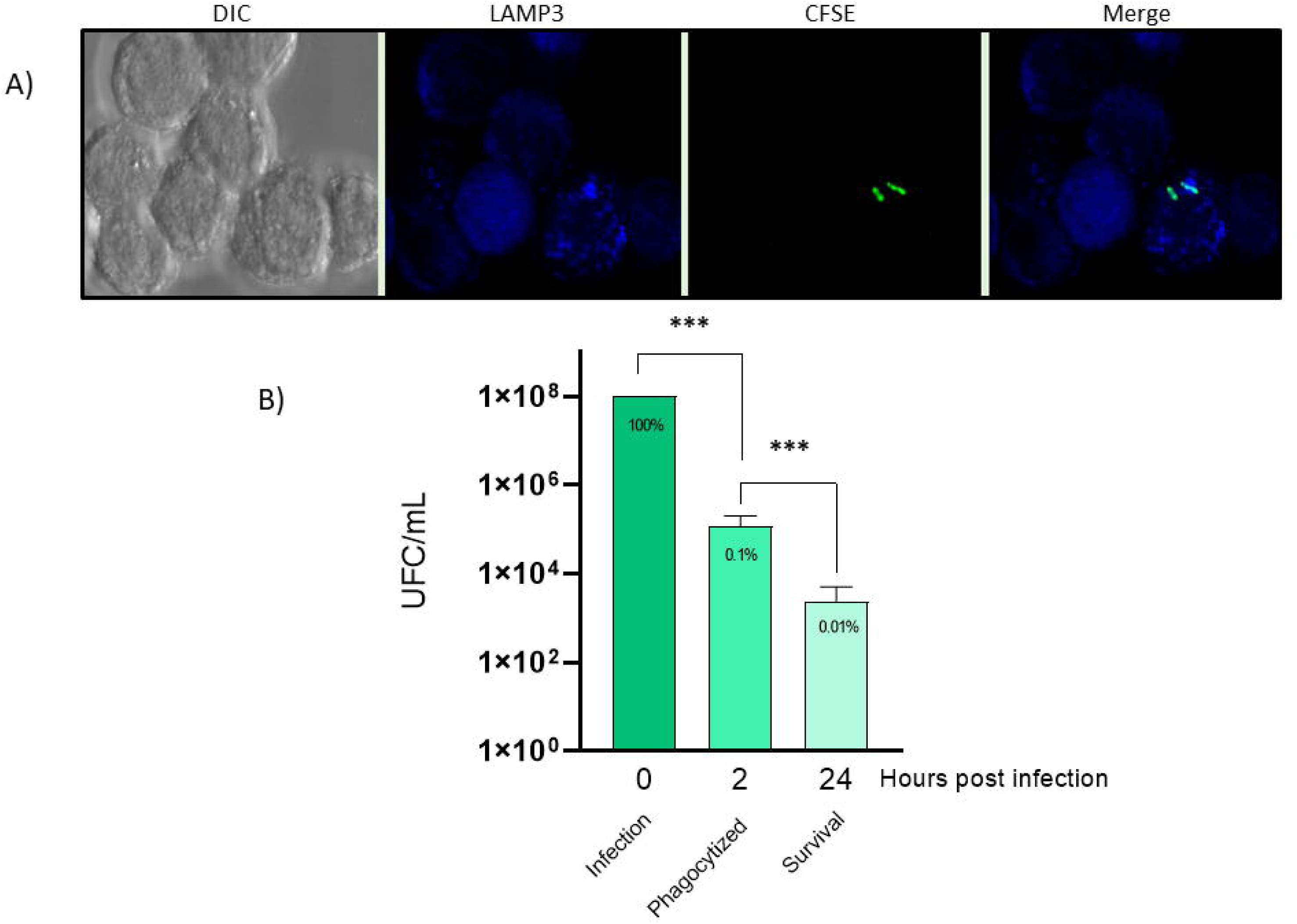
Heat map of gene expression. A) Ribosome assembly, B) Response to oxidative stress, C) OxyR regulon, D) SoxS regulon and E) SoxR regulon in EHEC O157:H7 phagocytized and *in vitro* stress. Heat maps depicting log_2_FC in the expression of EHEC genes at 2 h of incubation. Expression values are represented colorimetrically, with green representing upregulation, red downregulation and X as undetected. Data are the means from 3 biological replicates.

### Genes involved in the response to oxidative stress

*Escherichia coli* has two regulatory defense systems activated during oxidative stress conditions: the OxyR system [30] and the SoxRS system [30,31] which respond to hydrogen peroxide (H_2_O_2_) and redox-active compounds, respectively. Additionally, RpoS and Fur regulate several genes involved in response to oxidative stress [32].

The transcriptomic results for the phagocytized EHEC O157:H7 showed an upregulation of *oxyR, soxRS* and *rpoS* transcription. In contrast, downregulation of *fur* was seen (Fig 6B). OxyR is a constitutively expressed protein whose activation depends directly on H_2_O_2_ [30]. Subsequently, the oxidized OxyR initiates the transcription of several antioxidant genes, including *katG* (coding for hydroperoxidase I), *ahpCF* (coding for an alkyl hydroperoxide reductase), *dps* (coding for a nonspecific DNA binding protein) and *grxA* (coding for glutaredoxin I) [33].

Despite the overexpression of *oxyR,* there was no upregulation of these genes associated with oxidative stress by H_2_O_2_ in phagocytized EHEC O157:H7. Therefore, this would indicate that at 2 h of phagocytosis, the phagosome lumen still lacked H_2_O_2_ or had low concentrations of this stressor and was unable to activate the OxyR defense pathway.

In contrast, EHEC O157:H7 grown under H_2_O_2_ oxidative stress conditions also showed upregulation of *oxyR* followed by transcription of H_2_O_2_ oxidative stress-related genes (Fig 6C).

This finding demonstrates that *oxyR* transcription is insufficient to upregulate genes against oxidative stress, although the presence of H_2_O_2_ is necessary to activate OxyR and, therefore, develop the defense response.

The *soxRS* locus consists of two genes, *soxR* and *soxS*, with divergent transcription patterns. The SoxR protein is synthesized constitutively and becomes active in the presence of superoxide agents. Once oxidized, SoxR enhances the transcription of the *soxS*, whose product also acts as a transcriptional activator [34].

In this study, the *soxRS* gene displayed an upregulated profile in phagocytized EHEC O157:H7. However, the transcription pattern of the inducible genes forming part of the *soxRS* regulon was downregulated or remained unchanged. These genes include *acrAB* (an efflux pump), *fpr* (NADPH-ferredoxin reductase), *fur* (regulator involved with iron metabolism), *nfo* (DNA repair endonuclease IV), *fldAB* (flavodoxins), *sodA* (manganese-containing superoxide dismutase), and *zwf* (glucose-6-phosphate dehydrogenase) (Fig 6 D and E) [35–37].

The transcription patterns of these genes may be due to the lack of superoxide agents to activate SoxR in the lumen of the phagosome at 2 h. However, an alternative mechanism activates SoxR through nitric oxide (NO) [38]. The upregulation of *hcp* (Hydroxylamine reductase) a NO detoxification-associated enzyme [39] and *norR* (Anaerobic nitric oxide reductase transcription regulator NorR), [40], indicates the exposure of EHEC O157:H7 to macrophage-generated nitric oxide. The norR factor, which was found to be upregulated in phagocytized EHEC [41]. Furthermore, these nitric stress conditions could explain the activation of SoxR that generates an elevated fold-change of the *soxS* transcript observed in EHEC O157:H7 within the phagosome. EHEC O157:H7 *in vitro* stress conditions had no activated SoxR pattern, as these bacteria were not exposed to superoxide or nitric agents.

SoxR and SoxS also participate in the homeostasis response during harmful conditions [42]. For instance, SoxR directly activates the expression of metabolic genes involved in pathways of amino acid biosynthesis to counteract the potential inactivation caused by stress. The genes encoding the three susceptible enzymes AroF (DAHP synthase), TyrA (chorismate mutase/prephenate dehydrogenase), and MetE (homocysteine transmethylase) showed upregulation transcription levels in phagocytized EHEC O157:H7 (Fig 6D). Indeed, all these enzymes participate in the synthesis of amino acids (Hondorp and Matthews, 2004; Sobota et al., 2014); therefore, their expression prevents the inactivity produced by this stress.

On the other hand, SoxR and SoxS participate in the transcription of genes implicated in lipid biosynthesis in response to external stress. The LpxC enzyme (UDP-3-O-acyl-N-acetylglucosamine deacetylase), which is implicated in the catalysis of lipid A biosynthesis, was upregulated in phagocytized EHEC O157:H7. In addition, SoxR proved to be involved in the transcription of divalent ion transporter genes such as *zinT* and *znuABC* for Zn^2+^ and *mgtA* for Mg^2+^ transportation in phagocytized EHEC O157:H7. Several enzymes activated in stress conditions, such as TrxC, LpxC and MetE, depend on Zn^2+^ to act accurately [42]. Regarding Mg^2+^, this chemical element is used to activate sites of DNA polymerases, ATPases and kinases.

### Acid stress response

There is strong evidence that the EvgS/EvgA system is at the top of a pathway associated with acid and multidrug resistance in *E. coli* [2–10]. EvgS kinase sensor is stimulated by mild acidity (pH 5.5-5.7) and subsequent occurs the phosphorylation of the EvgA regulator [9–11]. EvgA activated regulates transcription of *safA-ydeO* operon, *emrK* and *yfdX* genes [43,44] We observed that these genes were upregulated in phagocytized EHEC O157:H7 with high foldchange values (Fig 7A). This would indicate an acid pH-respond activity of EvgA. In turn, *phoQ* and *phoP* were also upregulated. SafA activates in acid conditions the sensor kinase PhoQ, thereby connecting the EvgS/EvgA and PhoQ/PhoP two-component signaling systems [17, 18].

**Figure 7ABCDEF.**
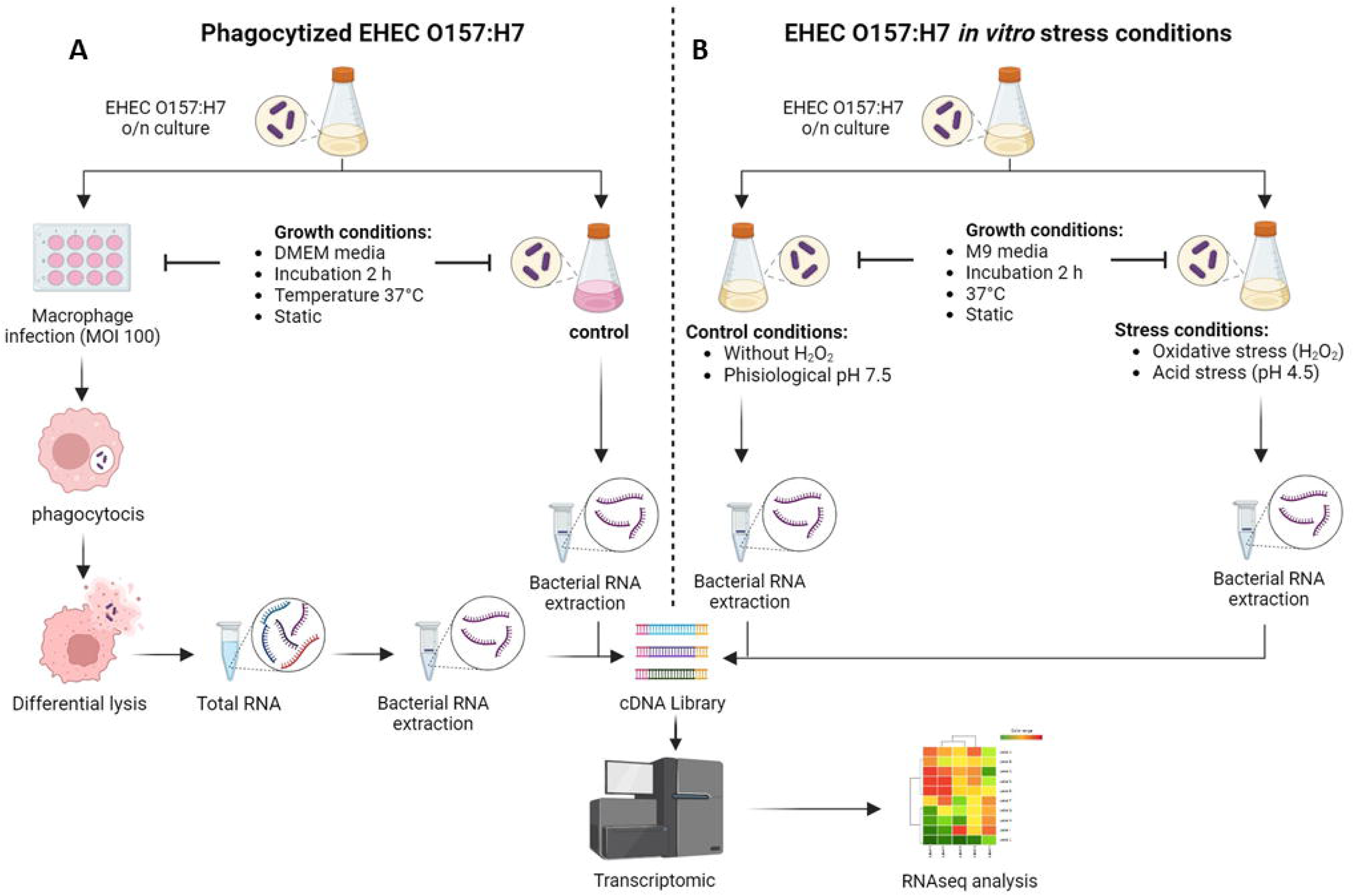
Heat map of gene expression. A) acidic stress, B) SOS response, C) TSS3, D) TSS6, E) Flagellar and F) Shiga toxin genes. Heat maps depicting log_2_FC in the expression of EHEC genes at 2 h of incubation. Expression values are represented colorimetrically, with green representing upregulation, red downregulation and X as undetected. Data are the means from 3 biological replicates.

Additionally, *E. coli* has five acid response systems (AR1-5) to resist extreme acidic conditions. In this regard, AR1 is a glucose-repressed or oxidative system activated by RpoS independently of the presence of external amino acids [45]. The AR2-5 systems, conversely, do rely on a specific extracellular amino acid and comprise an antiporter and a decarboxylase enzyme [46,47]. These systems confer acid resistance by utilizing glutamate, arginine, lysine, and ornithine as substrates for the decarboxylation reaction, which results in the consumption of intracellular protons with the consequent lower acidic internal pH [48].

All of the AR systems had a downregulated or unregulated profile in phagocytized EHEC O157:H7 (Fig 7A).

Therefore, these results suggest that the pH of the phagosome lumen reach a mild acidic condition after 2 h of phagocytosis, which indicates that the phagosome corresponds to an early stage. As the phagosome matures, it gradually decreases its internal pH to more extreme conditions to degrade the phagocytized pathogen. This acidification is achieved through the activity of proton pumps on the phagosome membrane, which actively transport hydrogen ions into the phagosome.

In contrast, EHEC O157:H7 *in vitro* stress conditions maintained a constant pH of 4.5 throughout the experiment. This finding demonstrates the expression of AR2-5 systems. In particular, the expression pattern of the primary genes related to AR2, such as *gadABC*, was overexpressed. Overexpression of *gadE*, one of the main regulators interacting in this network, was also evident. GadE is involved in the spatial and temporal regulation of gene expression [49]. HdeA and HdeB are two periplasmic chaperones transcribed as part of the GadE- dependent “acid fitness island” [50].

In certain *E. coli* strains, including pathogenic EHEC strains, the absence of these proteins can lead to a significant reduction, by two to three times, in the survival rates under low pH conditions [51]. This island also contains the *hdeD, yhiD* and *slp* genes, many of which have unknown functions. These components were upregulated in EHEC O157:H7 *in vitro*, thus demonstrating its involvement in the participation of the acid stress response.

AR2, on the other hand, presents an alternative circuit that can be activated by the expression of the *rpoS-gadX-gadY-gadW* transcripts during the stationary phase of growth [52]. Regarding AR4, the *cadA* and *cadB* genes showed an upregulated profile in EHEC exposed to stress *in vitro*, which indicates that this system would also be active under these conditions.

The activity of the AR4 system depends on two factors present in the assay: an acidic pH level and the presence of lysine in the minimum media. Decarboxylase CadA uses an external lysine to convert the amino acid to cadaverine under one proton consumption. The membrane-integrated one-component regulator CadC activates the expression of the *cadBA* operon, which is representative of the ToxR family [53].

Regarding AR3, also known as the Adi system, is induced under acidic pH conditions and an arginine-containing medium. AdiA is the arginine decarboxylase, whereas AdiC is the arginine/agmatine antiporter. These two genes were upregulated *in vitro*. This is also consistent with the increased expression of *adiY* since the product of this gene acts as a positive regulator of adiA transcription under low pH conditions [54].

Finally, the *speF* and *potE* genes, whose products are responsible for the activity of the AR5 system under acidic conditions, also showed an upregulated profile. PotE imports ornithine and exports putrescine from the membrane, while SpeF acts as an ornithine-decarboxylation enzyme to reduce cytoplasmic protons [55].

### SOS response

The regulation of the SOS response involves LexA and RecA as two pivot proteins. During regular growth, LexA serves as a transcriptional repressor for genes within the SOS regulon [56]. Under DNA damage conditions, RecA is activated as a co-protease to trigger the self-cleavage of LexA and other associated proteins [57]. Consequently, the levels of LexA start to decline and this decrease initiates the derepression of SOS genes. The SOS response includes excision repair, homologous recombination, translesion DNA replication and cell division arrest [58].

In phagocytized EHEC O157:H7, *uvrA*, *recA*, *ruvA, ruvB,* and *recN* displayed an upregulated profile (Fig 7B). The first gene induced during the excision of damaged nucleotides is *uvrA*, followed by *recA*. The response continues with the transcription of *ruvA, ruvB* and *recN*, which encode proteins involved in homologous recombination. The expression of these genes is associated with minor endogenous DNA damage [58]. Simultaneously, the *polB* (DNA polymerase II), *dinB* (DNA polymerase IV) and *sulA* (cell division inhibitor protein) genes also showed an upregulated pattern. This expression response takes place when environmental hostility to bacterial DNA persists. In addition, this sample showed elevated expression of the *umuD* and *umuC* genes. The *umu* operon is associated with the production of DNA polymerase V [59,60]. The induction of the transcription of these two genes occurs when the damage is more extensive.

This finding indicates that the DNA polymerase II and IV activity were not enough to repair the DNA damage. In contrast to other DNA polymerases, DNA polymerase V has a higher processivity but is more error-prone because of the lack of a proofreading activity. However, this induction allows it to repair the DNA under these conditions. Regarding the cell division, the decrease in gene transcription associated with cell replication and the expression of arrest cell division proteins indicate that phagocytized bacteria cannot divide under conditions of phagosome stress. Therefore, the bacteria persisted within the macrophage without replicating, as evidenced in the infection assays where no increase in CFU/ml was observed at 24 h.

Under *in vitro* stress conditions, EHEC O157:H7 upregulated the expression of *recA*, *uvrA* and *recN*, all of which are genes associated with an initial SOS response involved in minor DNA damage.

### Virulence factors

#### Type 3 and 6 secretion systems

T3SS is encoded in a pathogenicity island known as the locus of enterocyte effacement (LEE). This island comprises 41 open reading frames organized in five polycistronic operons known as LEE1–LEE5 [61,62]. Ler, the master transcriptional regulator encoded in the LEE1 operon, actively stimulates the expression of genes encoded from LEE2 to LEE5. The remaining genes in LEE1, as well as in LEE2 and LEE3, encode the fundamental structural components of T3SS. Additionally, LEE4 encodes supplementary T3SS structural components, translocators, and effector proteins. Finally, LEE5 encodes the crucial adhesin intimin (*eae*) and its corresponding “translocated intimin receptor” *tir*, which plays an indispensable role in establishing a close bond with host epithelial cells [63].

Both phagocytized EHEC O157:H7 and samples under *in vitro* stress conditions showed upregulated transcription patterns of the *ler* (master transcriptional regulator of the LEE operons), coincident with the downregulation of most ORFs associated with structural, translocator and effector proteins (Fig 7C). A regulatory pathway described in EHEC O157:H7 that senses intestinal metabolites is the EvgSA complex. This complex activates LEE gene expression by direct binding to the *ler* promoter [64]. The phagosome lumen lacks these metabolites and, as expected, the EvgSA complex transcription was downregulated after bacterium phagocytosis. The low levels of EvgSA resulted in the downregulation of Ler, with the subsequent repression of the T3SS virulence mechanism. Similarly, the EHEC O157:H7 under *in vitro* stress conditions did not recognize the signals as intestinal environment, as evidenced by the downregulation of *ler* and unchanged expression of *evgS* and *evgA*.

On the other hand, the EHEC O157:H7 strain also has another virulence-associated type 6 secretion system subtype 2 (T6SS-2). T6SS-2 has recently been associated with disease in a murine model linked to bacterium survival within the macrophage [65]. The T6SS-2 locus encodes minimal core components of 13 proteins in two operons. One operon contains TssjLM proteins that form the membrane complex: TssEFGHK, which comprises the base plate; the TssBC protein complex, which acts as a contractile sheath; and the ATPase ClpV, which is essential for recycling the sheath components [66]. The other operon contains Hcp, which forms the hexamers that make up the tail; VgrG, which constitutes a trimer and therefore associates with the tail to act as a spike; and some putative Rhs effectors with unknown functions [67].

Wan et al. (2017) have demonstrated that the mutant Δ*hns* (histone-like global repressor H-NS) EHEC O157:H7 EDL933 strain upregulated T6SS-2 gene transcription during macrophage phagocytosis [68]. They also observed an upregulation of *katN*, a characterized effector gene encoded outside the T6SS-2 operon. KatN exhibited a catalase activity when translocated by T6SS-2 to the phagosome and, in turn, this neutralized the macrophage ROS activity. However, our experiments revealed that the components of T6SS-2 remained unaltered or exhibited downregulation when bacteria were phagocytized or subjected to *in vitro* stress conditions (Fig 7D). Importantly, in the wild-type EHEC O157:H7 EDL933 strain, the authors detected no upregulation of T6SS-2 or its effectors. This finding would suggest that H-NS negatively regulates T6SS-2. In this study, *hns* was found to be downregulated in the phagocytized bacteria, whereas no changes in *hns* expression were observed in the bacteria exposed to *in vitro* stress.

#### Genes associated with motility

The biosynthesis and assembly of a flagellum involves over 50 genes organized into at least 14 operons [69,70]. These flagellar genes are classified into three hierarchical classes (Class I, II, and III) based on their specific timing and mode of expression [71]. The regulatory cascade of flagellar gene expression is tightly controlled by FlhDC (Class I), a protein that encodes the master flagellar regulator of Class II genes. FliA and FlgM Class II are responsible for synthesizing the hook-basal body structure and two regulatory proteins interacting within the Class III gene group. In addition, Class III proteins play a crucial role in synthesizing the entire flagellum and the chemotaxis system.

In this study, phagocytized EHEC O157:H7 showed a downregulated pattern of the Class I *flhC* and *fhlD* transcription; which avoided the subsequent cascade of class II and III gene expression. This finding indicates that phagocytized EHEC O157:H7 downregulates the expression of flagellar genes, with the consequent inhibition of flagellar biosynthesis (Fig 7E). Precise regulation of flagellar gene expression is essential to ensure resource conservation inside the phagosome to survive [72]. Additionally, the absence of flagellin avoids the triggering of an immune response by prompting the secretion of proinflammatory chemokines in intestinal epithelial cells [73].

In contrast, under *in vitro* stress conditions, the transcription of *flhA, flhB* and *flhC* were upregulated; therefore, flagellar biogenesis remained unaffected (Fig 7E). This result is reasonable since the ability to move using the flagellum provides bacteria with the significant advantage of being able to move towards favourable conditions and evade harmful environments.

#### Shiga toxin

Stx is recognized as the primary virulence factor of EHEC [74]. Its synthesis appears to be co-regulated through the activation of the integrated bacteriophage that encodes the toxin gene [75], a process linked to the induction of the bacterial SOS response, a universal reaction to DNA damage [76]. Given the observed upregulation of genes associated with the SOS response, we decided to assess the differential expression profile of Shiga toxin. In our experiments, we evaluated the differential expression of *stx2a* and *stx1a* in EHEC phagocytized by immune cells, as well as under *in vitro* stress conditions. The results showed a marked overexpression of *stx2a* in phagocytized EHEC, suggesting that this gene is particularly sensitive to phagocytosis, which could enhance virulence during host interactions (Fig 7F). On the other hand, *stx1* was downregulated under these same conditions (Fig 7F). However, under *in vitro* stress conditions, both *stx1* was upregulated, indicating that the mechanisms controlling Shiga toxin gene expression may vary depending on the type of stress the bacterium faces (Fig 7F). These results support the idea that the regulation of Stx expression is influenced by both internal and external signals, which may have implications for the severity of the disease in human infections.

## Conclusion

Based on the observed results, the defense strategy of an early murine phagosome involves triggering the bacterial SOS response, lipid biosynthesis and NO detoxification enzymes. This response is induced in reaction to DNA, membrane damage, NO stress and mildly pH conditions. Several enzymes activated in these stress conditions, such as DNA polymerases, ATPases and kinases, depend on divalent ions whose transporters were also upregulated. Subsequently, to survive, the bacterium conserves energy by downregulating flagellar biosynthesis, components of the T3SS and T6SS virulence mechanisms. In this context, the acidic pH, reactive nitrogen species, and oxidative stress present in the phagosome create significant challenges for bacterial persistence. By upregulating *stx2a*, EHEC may enhance its virulence, potentially disrupting host cell processes, weakening immune defenses, and facilitating escape from the immune system. This underscores the importance of Shiga toxin as a key factor in the bacterial response to intracellular stress, contributing to the pathogen’s ability to persist and cause disease. At the same time, the bacterium increases the ribosomal levels to react more effectively against adverse conditions to enhance its response to improve environmental conditions.

Under more extreme stress conditions, EHEC O157:H7 upregulated the expression of genes to overcome those conditions. For example, in an acidic environment (pH 4.5), the bacterium upregulated the acid stress response pathways AR2-5, whereas, upon high concentrations of hydrogen peroxide, it transcribed higher levels of genes of the OxyR pathway and subsequently H_2_O_2_ oxidative stress-related genes.

Altogether, the RNAseq analysis provided a deeper understanding of the mechanisms involved in the persistence of the EHEC O157:H7 strain within the phagosome.

## Supporting information

RNA-seq

## Acknowledgments

We thank Jinlong Bei, who kindly provided us with the necessary resources to carry out the experiences.

## Author Contributions

**Conceptualization:** Nahuel A. Riviere, Jinlong Bei, Ángel Adrián Cataldi and Mariano Larzábal.

**Data curation:** Wanderson Marques Da Silva, Ángel Adrián Cataldi, Lin Jiang, Xiuju Wu, Ke Xing, María Carolina Casabonne, Lu Guo and Libia Yael Smith.

**Formal Analysis:** Nahuel A. Riviere, Ke Xing and Mariano Larzábal.

**Funding acquisition:** Jinlong Bei, Ángel Adrián Cataldi and Mariano Larzábal.

**Investigation:** Nahuel A. Riviere, Ke Xing and Mariano Larzábal.

**Methodology:** Nahuel A. Riviere, Ke Xing and Mariano Larzábal.

**Project administration:** Nahuel A. Riviere, Jinlong Bei, Ángel Adrián Cataldi and Mariano Larzábal.

**Resources:** Lin Jiang, Xiuju Wu, María Carolina Casabonne, Lu Guo, Libia Yael Smith and Wanderson Marques Da Silva.

**Software:** Nahuel A. Riviere, Ke Xing and Wanderson Marques Da Silva.

**Supervision:** Jinlong Bei, Ángel Adrián Cataldi and Mariano Larzábal.

**Validation:** Nahuel A. Riviere, Ke Xing and Mariano Larzábal.

**Visualization:** Nahuel A. Riviere and Mariano Larzábal.

**Writing – original draft:** Nahuel A. Riviere and Mariano Larzábal.

**Writing – review & editing:** Nahuel A. Riviere, Jinlong Bei, Ángel Adrián Cataldi, Ke Xing and Mariano Larzábal.

## Supporting information

S1 Fig. Quality control of RNA samples from phagocytized EHEC O157:H7 exposed to in vitro stress.

S2 Fig. Principal component analysis.

S3 Table. Quantification of the RNA samples obtained for the different conditions.

S4 List of genes obtained by RNA-Seq analysis in phagocytosed EHEC and EHEC exposed to in vitro stress. The tag locus of each gene, the fold change corresponding to the differential expression under the experimental conditions, and the associated p-value are included.

