## Supplementary figures and images for "Global expression profile of Enterohemorrhagic *Escherichia coli* O157:H7 in phagosome of murine macrophages"

### Fig S1

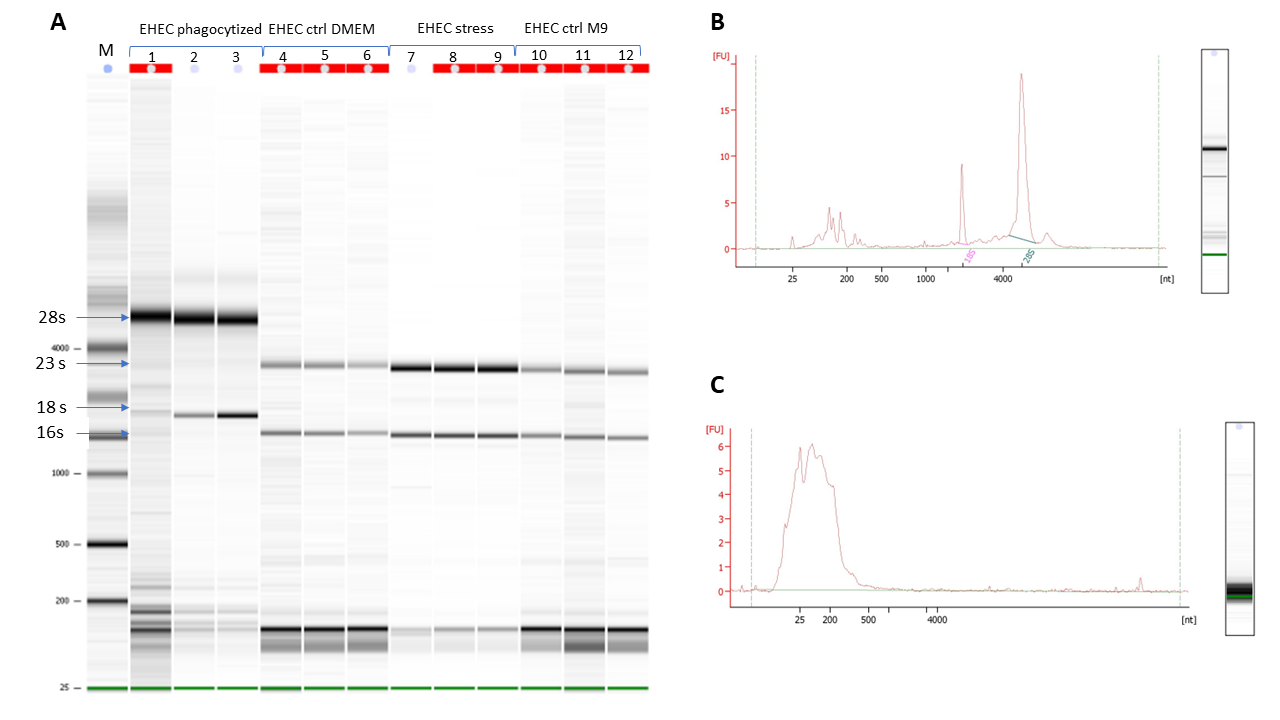

### Fig S2

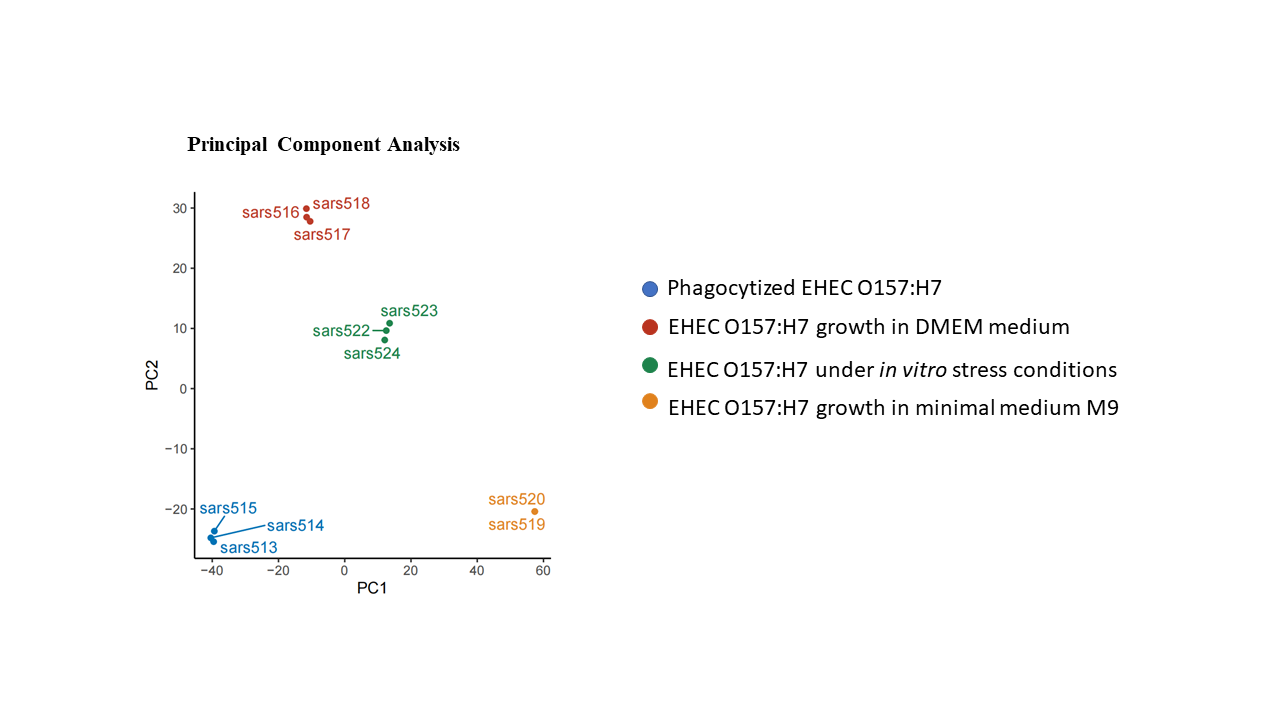

### Table S3

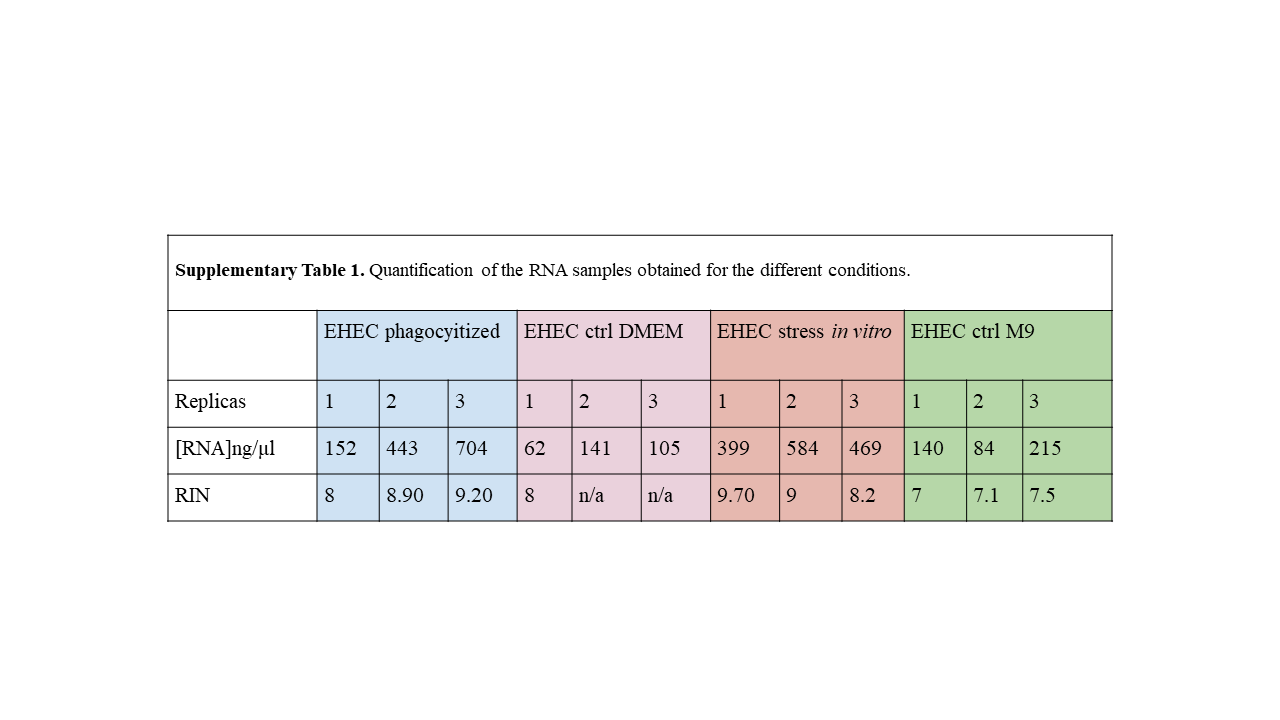
