## Supplementary material for "Global expression profile of Enterohemorrhagic *Escherichia coli* O157:H7 in phagosome of murine macrophages": Table S1

| **Supplementary Table 1.** Quantification of the RNA samples obtained for the different conditions. | | | | | | | | | | | | |
| --- | --- | --- | --- | --- | --- | --- | --- | --- | --- | --- | --- | --- |
|  | **EHEC phagocyitized** | | | **EHEC ctrl DMEM** | | | **EHEC stress *in vitro*** | | | **EHEC ctrl M9** | | |
| **Replicas** | 1 | 2 | 3 | 1 | 2 | 3 | 1 | 2 | 3 | 1 | 2 | 3 |
| **[RNA]ng/μl** | 152 | 443 | 704 | 62 | 141 | 105 | 399 | 584 | 469 | 140 | 84 | 215 |
| **RIN** | 8 | 8.90 | 9.20 | 8 | n/a | n/a | 9.70 | 9 | 8.2 | 7 | 7.1 | 7.5 |

**Supplementary Table 1**. RNA quantification from samples obtained under different experimental conditions: phagocytized EHEC and EHEC under *in vitro* stress, along with their respective controls (EHEC control DMEM and EHEC control M9). Quantification was performed by capillary electrophoresis using the Agilent 2100 system, with RNA concentration in ng/μl and the RNA integrity number (RIN) reported. The RIN value ranges from 1 to 10, where an RIN of 10 indicates completely intact RNA, while an RIN of 1 reflects total degradation. n/a: not assigned.
